# Mucociliary Clearance Augmenting Drugs Block SARS-Cov-2 Replication in Human Airway Epithelial Cells

**DOI:** 10.1101/2023.01.30.526308

**Authors:** Javier Campos-Gomez, Courtney Fernandez Petty, Marina Mazur, Liping Tang, George M. Solomon, Reny Joseph, Qian Li, Jacelyn E. Peabody Lever, Shah Hussain, Kevin Harrod, Ezinwanne Onuoha, Harrison Kim, Steven M. Rowe

**Affiliations:** Department of Medicine, University of Alabama at Birmingham, Birmingham, Alabama; Gregory Fleming James Cystic Fibrosis Research Center, University of Alabama at Birmingham, Birmingham, Alabama; Medical Scientist Training Program, Heersink School of Medicine, University of Alabama at Birmingham, Birmingham, Alabama; Department of Anesthesiology and Perioperative Medicine, University of Alabama at Birmingham, Birmingham, Alabama; Biomedical Engineering, University of Alabama at Birmingham, Birmingham, Alabama; Department of Radiology, University of Alabama at Birmingham, Birmingham, Alabama

**Keywords:** Mucociliary transport, mucoactive drug, SARS-CoV-2, cilia movement, antioxidant

## Abstract

The coronavirus disease (COVID-19) pandemic, caused by SARS-CoV-2 coronavirus, is devastatingly impacting human health. A prominent component of COVID-19 is the infection and destruction of the ciliated respiratory cells, which perpetuates dissemination and disrupts protective mucociliary transport (MCT) function, an innate defense of the respiratory tract. Thus, drugs that augment MCT could improve barrier function of the airway epithelium, reduce viral replication and, ultimately, COVID-19 outcomes. We tested five agents known to increase MCT through distinct mechanisms for activity against SARS-CoV-2 infection using a model of human respiratory epithelial cells terminally differentiated in an air/liquid interphase. Three of the five mucoactive compounds tested showed significant inhibitory activity against SARS-CoV-2 replication. An archetype mucoactive agent, ARINA-1, blocked viral replication and therefore epithelial cell injury, thus, it was further studied using biochemical, genetic and biophysical methods to ascertain mechanism of action via improvement of MCT. ARINA-1 antiviral activity was dependent on enhancing the MCT cellular response, since terminal differentiation, intact ciliary expression and motion was required for ARINA-1-mediated anti-SARS-CoV2 protection. Ultimately, we showed that improvement of cilia movement was caused by ARINA-1-mediated regulation of the redox state of the intracellular environment, which benefited MCT. Our study indicates that Intact MCT reduces SARS-CoV-2 infection, and its pharmacologic activation may be effective as an anti-COVID-19 treatment.

## Introduction

The SARS-CoV-2 coronavirus is the causal agent of the coronavirus disease (COVID-19) pandemic, posing a devastating impact on human health and, consequentially, in almost every aspect of society. Although vaccination has improved the incidence and severity of the disease, severe respiratory illness affects a substantial portion of the population and has been worsened by the emergence of more contagious variants of SARS-CoV-2. Still, specific immunocompromised populations, such as those suffering cancer, HIV or organ transplant recipients cannot sustain a good vaccine response (1, 2), and even people with adequate responses may still succumb to a continuum of long-term COVID-19 manifestations (3) Antiviral treatments such as monoclonal antibodies (e.g., sotrovimab; bebtelovimab) and small molecule antivirals (e.g., remdesivir, paxlovid, molnupiravir) have reduced disease severity (3). However, many of those monoclonals are not effective for more recent emerged Omicron lineages, a trend that is expected for other emerging variants (4). Additionally, corticosteroid treatment has reduced mortality in individuals with severe forms of the disease (5). Nevertheless, global morbidity and mortality remain unacceptably high in both the general population and in people with one or more risk factors for severe infection, with global daily incidence and death counts at 640,182 and 2,436, respectively on December 28, 2022 (6). Therefore, therapeutic interventions are strongly needed as alternative tools when vaccination or available treatments fail or are not an option.

A hallmark of COVID-19 respiratory disease is abnormal or absent ciliation of the respiratory epithelium, which disrupts the protection of the mucociliary transport (MCT) apparatus, an innate defense of the lung (7, 8). The reduction or loss of MCT by SARS-CoV-2 is expected to have consequential effects on COVID-19 illness, including accelerating or worsening descending infection of the airways, prolonged illness due to mucus plugging, and risk for secondary infections. As viral particles accumulate and spread within the deep lung, they contribute to mucus plugging, a prominent finding in severe cases of COVID-19. Mucus plugging increases the risk of secondary infections, which are commonly reported in influenza and COVID-19 (9, 10). Together, disrupted MCT and mucus plugging can lead to dysregulated inflammatory processes and lung damage, as well as long-term complications from COVID-19 (11). Considering these findings, re-establishing or augmenting MCT may have a direct effect on viral transmission to ciliated airway epithelial cells and may, in turn, have long-term implications on the prognosis and development of pulmonary complications from COVID-19, including long COVID, that could be precipitated by secondary consequences of altered mucus clearance.

We hypothesized that drugs that can augment MCT could block or reduce viral infection or cell damage to the respiratory epithelium. To evaluate this hypothesis, we tested a panel of agents that directly or indirectly improve MCT for their activity against SARS-CoV-2 *in vitro*. To assess the functional and mechanistic aspects of these drugs, we used terminally differentiated human bronchial epithelial cells (HBEC) grown in an air-liquid interface (ALI), an established model of SARS-CoV-2 infection. ALI cultures closely recapitulate key characteristics of the airway *in vivo*. Therefore, this model can be used to interrogate the mechanisms of tissue damage and cell death, as well as aspects of host response to infection (12, 13). Importantly, human airway cultures grown in an ALI system is a physiologically relevant model for investigating anti-COVID-19 interventions because the cells express angiotensin-converting enzyme 2 (ACE2), the host cell receptor of SARS-CoV-2, and the serine protease TMPRSS2 that is required for the viral glycoprotein (protein S) priming (14, 15).

We initially tested a panel of five pharmacotherapies that augment MCT but have distinct mechanisms of action, as proven by previous evidence of bioactivity in respiratory disorders. Three of the tested mucoactive compounds showed a significant inhibitory activity against SARS-CoV-2 replication in the HBE-ALI antiviral assay. Based on bioactivity, we then examined the agent with the greatest efficacy, ARINA-1, in more detail. This inhaled investigational drug is a combination of glutathione, ascorbic acid, and sodium bicarbonate that has also been shown to improve MCT in HBEC (16, 17). Here we showed that ARINA-1 antiviral activity is dependent on improvement of MCT, since the drug is not active in the absence of terminal differentiation or ciliary motility. Our findings suggest that MCT-activating drugs may be effective as anti-COVID-19 treatment, deserving further exploration. Our study emphasizes the importance of a properly functioning mucociliary escalator to protect the airways from SARS-CoV-2 or other viral pathogens that replicate within respiratory epithelial cells.

## Materials and Methods

### Procurement and growth of human primary airway epithelial cells

The UAB Institutional Review Board approved the use of human cells and tissues. Primary HBEC were derived from lung explants after written informed consent was obtained from donor subjects using methods described previously (18, 19) [see details in data supplement]. Briefly, tissues were debrided immediately after surgical resection, washed twice in Minimum Essential Media with 0.5 mg/ml DTT (Sigma-Aldrich, St. Louis, MO) and 25 U/ml DNAse I (Roche, Basel, Switzerland), and then placed in dissociation media containing MEM, 2.5 U/ml DNAse I, 100 mg/ml ceftazidime, 80 mg/ml tobramycin, 1.25 mg/ml amphotericin B, and 4.4 U/ml pronase (Sigma-Aldrich) for 24–36 h at 4°C. Loosened airway epithelial cells were then expanded using BEGM medium (LONZA, Basel, Switzerland) supplemented with an additional 10 nM all-trans-retinoic acid (Sigma-Aldrich) that was exchanged every 24 h. Following expansion, first or second passage cells at 80–90% confluency were dissociated and seeded at a density of 0.5 × 10^5^ on Costar® Transwell 24-well filter inserts (cat. # 3470, Corning Inc.) after coating with NIH 3T3 fibroblast conditioned media. Cells were differentiated at ALI for at least five weeks, at least otherwise is specified, using medium PneumaCult−/heparin/hydrocortisone (Stemcell− Technologies) plus pen/strep antibiotics (exchanged three times a week) before further use. Cells from five different donors were used in this study: WT128, WT148, WT152, WT158 and WT210 cell lines.

Nasal cells from two healthy and two PCD patients [patient 1 genotype: *CCDC39* c.2586+1G>A (splice donor) heterozygous pathogenic and *CCDC39* c.830_831del (p.Thr277Argfs*3) heterozygous pathogenic; patient 2: *DNAI1* exon 5, c.370C>T (p.Arg124Cys) heterozygous pathogenic and intron 1, c.48+2dup heterozygous pathogenic] were obtained from nasal brushes after written informed consent was obtained from donors and grown as co-culture with 3T3 J2 cells. When the cells become 80-90% confluent the cells were detached using 0.05% trypsin. A total of 1.25 × 10^5^ cells were plated in FNC coated Costar® Transwell 24-well filter inserts (cat. # 3470, Corning Inc.). After three days medium from the apical compartments was removed and differentiation medium (Pneumacult ALI maintenance medium) maintained at the basolateral side only. Medium was changed every other day until terminal differentiation (two and a half to three weeks) and then used for the experiments.

### Culture of 16HBE at the ALI

16HBEC14o-were grown using EMEM medium (ATCC, cat. # 30-2003−) supplemented with 10% FBS and were dissociated at 80–90% confluence and seeded at a density of 0.5 × 10^5^ on Costar® Transwell 24-well filter inserts (cat. # 3470, Corning Inc.) after coating with FNC (Athena ES 0407H) and using EMEM medium. Upon reaching confluence, the medium was removed from the apical side, and the cells were grown at the ALI for no longer than two weeks to avoid differentiation.

### Compounds used in this study and their sources

The compounds used throughout this study and their sources (in parentheses) were the following: ARINA-1 (kindly donated by Renovion, Inc.); camostat mesylate (Millipore Sigma Cas # 59721-29-8); poly-N (acetyl, arginyl) glucosamine (PAAG, kindly donated by Synedgen, Inc.); allopurinol (TCI, Cat. # A0907); ascorbic acid (LETCO Medical, Cat. # 684471); reduced glutathione (Sigma, Cat. # G6013); sodium bicarbonate (LETCO Medical, Cat. # 685940); N-acetylcysteine (EMD Millipore, Cat. # 106425); L-; BAPTA/AM (EMD Millipore, Cat. # 196419); sulforaphane (Medkoo Biosciences Inc. Cat. # 202713250MG). Hydrosoluble and hydrophobic compounds were dissolved in sterile milli-Q water and dimethyl sulfoxide (DMSO), respectively. They were used at the indicated concentration depending on the experiment.

### HBEC/ALI-based antiviral assay

All the antiviral assays performed using the following method were done using fully differentiated primary HBEC from several human donors, unless stated otherwise. For the antiviral assay, the HBEC were washed two times to eliminate the mucus excess by adding 100 µl of PBS1X, incubating for 30 min at 37°C under 5% CO_2_, and aspirating the liquid without touching the cells. Subsequently, the vehicle (DMSO or saline, depending on the compound used) and test compounds were added at the specified concentrations to the basolateral or apical side of the cell layers as desired, and incubated for one hour at 37°C in a 5% CO_2_ incubator (only one microliter of the compounds tested apically was added per well). Cells were exposed for an additional hour to SARS-CoV-2 (Washington strain WA-1/USA ancestral clade A, obtained from BEI Resources) at an MOI=0.24. That is, a 30 µl aliquot of a viral stock at 1.37 × 10^7^ infectious virus/mL was added to each well, which contained an average of 1.7 × 10^6^ HBEC). Excess unattached viral particles were aspirated from the filters and washed with 100 µl of PBS. Since this washing step removed the apical compounds, they were added again at this point (1 µL). Control cells treated with vehicle and exposed to the virus for one hour were scraped from the transwell filters, collected in 100 µl of PBS, and frozen at -80°C for further analysis. These served as our t=0 baseline virus controls, i.e., initial virus amount attached or internalized before viral replication started. The rest of the filters, treated with either the vehicle or the test compounds, were incubated at 37°C under 5% CO_2_ for 72 hours. During this time, 1 uL of the apically tested compounds were added again at 24 and 48 h. That is, apical compounds were added four times: one hour before infecting the cells (which was washed out at the step of excess virus removal), then three times at 0, 24 and 48 h post infection (which were never removed from cell layer surface), unless stated otherwise. Compounds tested basolaterally were dissolved into the medium underneath the transwell filters one hour before virus exposure and kept for the duration of the experiment (i.e. the medium was never changed). At the end of the 72-h incubation period of the assay, cells were scraped and collected in 100 µl of PBS from each filter and frozen at -80°C for further analysis of the viral load together with the baseline controls collected at t=0.

### Measurement of viral load by RT-qPCR

The effect of the drugs on blocking SARS-CoV-2 infection of HBE was assessed by measuring the viral load through the quantification ofby quantifying the SARS-CoV-2 genomic RNA copy number by RT-qPCR. Total RNA was purified from all the samples, including the baseline controls collected at t=0, using the QIAamp Viral RNA Mini Kit (QiagenCat. # 52906). Purified RNA samples were diluted 1/50, and 5 µl of each dilution used to perform RT-qPCR to determine viral load (viral RNA copy number). The RT-qPCR was performed using the 2019-nCov CDC EUA kit (probe and oligos, IDT cat. # 10006776) and Promega® GoTaq® 1 Step RT PCR (Cat. # A6020) in a QuantStudio 3 thermocycler (Applied Biosystems) and following the recommendations of the CDC (https://www.cdc.gov/coronavirus/2019-ncov/lab/virus-requests.html). The standard curve of the assay was done using the 2019-nCoV_N_Positive Control plasmid (Cat. # 10006625, IDT). This plasmid was linearized with ScaI restriction enzyme to improve the dynamic range and assay-to-assay reproducibility of the standard curve. After cleaning the plasmid digestion using the Monarch DNA Cleanup Kit (NEB, #T1030), the DNA concentration of the linearized plasmid was measured to determine the number of DNA molecules in the stock solution, which was calculated using SnapGene software version 5.0.8. Six concentrations were prepared at 10^4^, 10^5^, 10^6^, 10^7^, 10^8^ and 10^9^ DNA molecules/mL for the standard curve, which was included in every PCR amplification of the antiviral assays. To avoid variability, enough of each point solution was prepared for all the assays performed in this study. Thus, PCR efficiency was consistent between PCR amplification (between 95 and 105% efficiency).

### Measurement of infectious virus by TCID_50_

ARINA-1-treated or untreated primary fully differentiated HBEC cells were infected with SARS-CoV-2 as described above for the HBEC/ALI-based antiviral assay. After 72 h of incubation the apical side of the cell layers were washed with 100 µl of PBS 1X to collect the virus particles. Collected virus suspensions were serially diluted (5-fold series) in EMEM medium (ATCC, cat. # 30-2003−) in sextuplicate and then 30 μL of viral dilution were added to a 30 μL volume of freshly plated Vero E6 cells (ATCC, cat. # CRL-1586^™^) at 5,000 cells/well in 384-well white plate (Corning, cat. # 3570). The plates were incubated at 37°C under 5% CO_2_ for 144 hour and virus cytopathic effect was measured using the CellTiter-Glo^®^ 2.0 Cell Viability Assay (Promega, Cat. # G9241) following the manufacturer instructions. The luminescence values were used to calculate the TCID50 of each virus sample using the Viral ToxGlo− Assay (https://www.promega.com/resources/protocols/technical-manuals/101/viral-toxglo-assay-protocol/).

### Cytotoxicity of compounds

The cytotoxicity of all compounds (except ARINA-1 that interferes with the assay) was measured using the CytoTox 96^®^ Non-Radioactive Cytotoxicity Assay (Promega, Cat. # G1780), which measures LDH enzyme release. ARINA-1 cytotoxicity was measured using the CellTiter-Glo^®^ 2.0 Cell Viability Assay (Promega, Cat. # G9241), which measures intracellular ATP. All assays were done in 16HBE cells cultured at the ALI in EMEM medium as described above. The cells were exposed to the compounds either basolaterally or apically, depending on their mechanism of action for 72 hours at the highest concentrations used in the antiviral assays. For the LDH kit the basolateral medium was directly assayed following the manufacturer instructions. For the ATP kit, the cells were scraped from the transwell filters and recovered in 100 µl of PBS 1X, transferred to a white opaque 96-well plate, then mixed with 100 µl of CellTiter-Glo reagents, and continued following manufacturer instructions. Toxicity of the compounds were reported as percent of the maximum toxicity of control cells that were treated with a lysis reagent supplied by the kits.

### Assessment of SARS-CoV-2 cytopathic effect through immunohistochemistry

ARINA-1 cytoprotective activity of the HBEC was evaluated through immunohistochemistry using cilia and apoptosis markers (acetylated β-tubulin and caspase-3, respectively). The unstained slides of 5 µm paraffin sections of the HBEC were baked overnight at 60°C then deparaffinized by dissolving with three changes of xylene and hydrated using graded concentrations of ethanol to deionized water. The tissue sections were subjected to antigen retrieval by 0.01 M sodium citrate buffer (pH 6) in a pressure cooker for 5 min (buffer preheated). Following antigen retrieval, all sections were washed gently in deionized water, then transferred into 0.05 M Tris-based solution in 0.15 M NaCl with 0.1% v/v Triton-X-100, pH 7.6 (TBST). Endogenous peroxidase was blocked with 3% hydrogen peroxide for 15 min. To reduce further nonspecific background staining, slides were incubated with 5% normal goat serum (Sigma, G9023) for 45 min at RT. Slides were then incubated at 4°C overnight with the corresponding primary antibody (anti-acetylated tubulin, Sigma T7451, mouse monoclonal, 1/60,000 dilution; anti-cleaved caspase 3, Cell Signaling 9579, rabbit monoclonal, 1/500 dilution; or anti-SARS-CoV-2 glycoprotein S, Invitrogen PA141165, rabbit polyclonal, 1/500 dilution). After washing with TBST, sections corresponding to tubulin staining were incubated with a goat anti-mouse IgG secondary antibody conjugated with HRP (Abcam ab6789, 1:500), while sections of caspase and protein S staining were incubated with goat anti-rabbit IgG secondary antibody conjugated with HRP (Abcam ab6721, 1:1,000). ImmPACT DAB Peroxidase (HRP) Substrate Kit (Vector Laboratories SK4105) was used as the chromogen and hematoxylin (no. 7221, Richard-Allen Scientific, Kalamazoo, MI) as the counterstain. To validate the specificity of the antibodies, control slides were used following the protocol described above, but at the step of primary antibody incubation PBS buffer was used instead of the primary antibody.

### Cilia length measurement

Image processing and analysis were implemented using ImageJ v1.53a (Bethesda, MD). An RGB histologic photograph was opened, and each cilium was traced using the draw (pencil) tool from the ciliated columnar epithelial cell surface to the tip of the cilium. After tracing all cilia in the photograph, the data type was converted from RGB to 8 bits, and the cilia images were segmented using a thresholding technique. Then the binary images of cilia were skeletonized, and the total number of cilia and their lengths were measured using the “Analyze Skeleton 2D/3D” function.

### Testing direct antiviral activity of ARINA-1 on SARS-CoV-2 virion

A 100 µl volume of ARINA-1 or saline was added to 100 µl of SARS-CoV-2 suspension (5.52 × 10^8^ viral particles/mL) and incubated for 1 hour at 100 µl 37°C. After incubation, the mixtures were added to 3K MWCO 0.5 mL filter (Pierce, Cat. No. 88512) and centrifuged at maximum speed to concentrate the virus 10 times (to a volume of 20 µl). Then, 300 µl of PBS 1X were added to the filter, and the virus was concentrated again by centrifugation to 20 µl volume. This PBS wash was repeated two times to eliminate the rest of the ARINA-1 components, and the recovered viral suspension was adjusted to the original volume of 100 µl to keep the same viral concentration as the original suspension. A 30 µl volume of this processed viral suspension was used to infect Vero E6 cells plated at the confluence in a 96 well plate or HBEC in ALI filters. The viral load was measured by qRT-PCR as described above. Controls were run in parallel using saline instead of ARINA-1.

### μOCT Image acquisition and analysis

Measurements of functional microanatomic parameters in cultured primary HBEC were performed using μOCT as previously described (20, 21). Specifically, MCT rate was determined using the time elapsed, and distance traveled of native particulates in the mucus over multiple frames. Four videos were randomly acquired at standard distances from the center of the wells (mid-way between the center and the border circle) for each well. MCT rates were normalized against the MCT average of baseline controls (uninfected cells treated with saline).

## Results

### HBEC/ALI-based antiviral assay validation

The antiviral assays were performed as depicted in a simplified scheme of the assay (Fig. 1A) and further detailed in the Methods section. To validate the assay, camostat mesylate was utilized because it has the dual action of blocking SARS-CoV-2 entry through TMPRSS-2 mediated priming of viral S protein (22) in addition to mucoactive properties. The mucoactive effect of camostat mesylate is via the inhibition of the epithelium sodium channel (ENaC)-activating protease, which enhances MCT by reducing ENaC mediated fluid resorption(23). As anticipated, camostat mesylate inhibited the replication of SARS-CoV-2 in HBEC in a dose-dependent manner (Fig. 1B). Camostat mesylate inhibited SARS-CoV-2 infection greater than 99% at concentrations higher than 16.62 µM compared with the DMSO vehicle (Fig. 1C). The compound was not cytotoxic at the maximum concentration tested in the antiviral assays as measured by lactate dehydrogenase (LDH) release (125 µM, Fig. S1 from data supplement), ruling out the observed decrease in viral load was due to cell death caused by the drug. These results validated the antiviral assay in our experimental conditions.

**Fig. 1.**
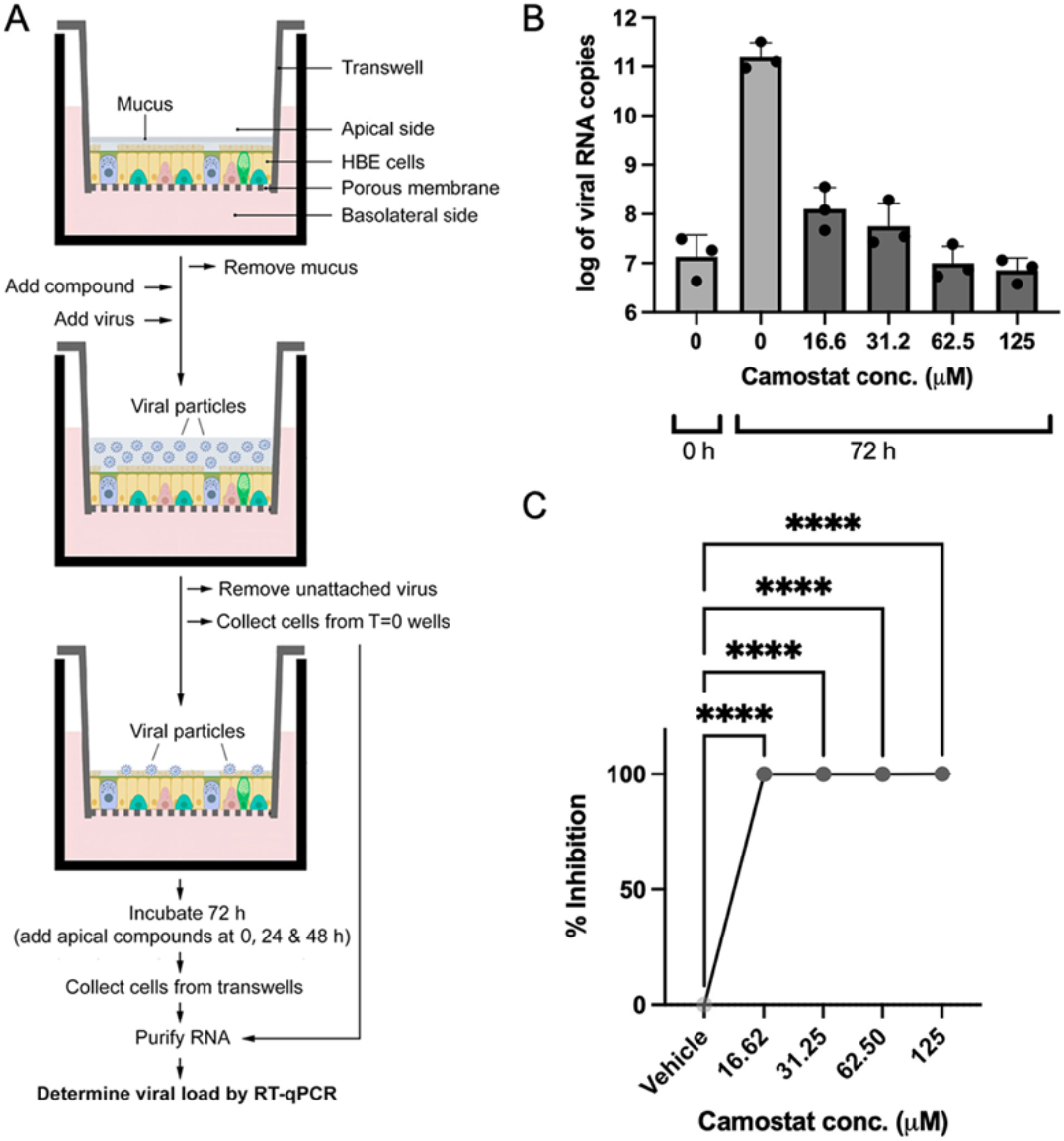
HBEC-ALI-based antiviral assay and validation. A) Schematic representation of the antiviral assay. Differentiated HBE cells grown at the ALI on transwells microporous filters are washed with PBS to remove mucus. Test compounds are added basolaterally (camostat, ivacaftor) or apically (PAAG, HA, ARINA-1). After one hour of incubation, SARS-CoV-2 virus is added at an MOI=0.24. Unattached viral particles are removed after an hour of incubation. At this point, the transwell data corresponding to time zero (before viral replication) was collected. The remainder of the transwells are incubated for 72 h with additional administrations of the apically delivered compounds (PAAG, HA, ARINA-1) at T=0, 24 and 48 h. Cells from each time point (0 and 72 h) are collected from the filters and used for RNA purification. Finally, the collected RNA is used to measure the viral load by RT-qPCR. B) Validation of the assay using camostat mesylate compound. A graph of the viral copy number measured by the RT-qPCR is shown for a representative experiment with three filter replicates per condition. Camostat and vehicle control were added basolaterally. DMSO was used as the diluent for camostat and the vehicle control. Data was logarithmically transformed. C) Data from two independent experiments, as the one shown in B, were converted to percent of viral inhibition, averaged (3 replicates/experiment) and compared using an ordinary one-way ANOVA (****p<0.0001).

### Effect of mucoactive compounds on SARS-CoV-2 infection of HBEC

We evaluated four additional mucociliary active drugs for potential antiviral activity: ivacaftor, poly-N (acetyl, arginyl) glucosamine (PAAG), high molecular weight hyaluronic acid (HA), and ARINA-1. Ivacaftor is a potentiator of the chloride ion transporter cystic fibrosis transmembrane conductance regulator (CFTR) that works by improving the function of the CFTR protein in cystic fibrosis (CF) patients with a gating defect 9,20. By restoring the flow of chloride ions through the cell membrane via CFTR, ivacaftor improves MCT, which reduces CF symptoms, such as the build-up of thick mucus in the lungs (24, 25). We and others have shown it has some bioactivity in wild type epithelia (26). The synthetic glycopolymer PAAG decreases the viscoelasticity of mucus in the airway and has been shown to be effective at reducing mucus viscosity and restoring MCT in CF cell culture models (27). The biopolymer HA, particularly its high molecular weight form, is a constituent of the extracellular matrix of the lungs with anti-inflammatory and water-retaining properties. Thus, HA plays a significant role in regulating fluid balance in the lung and airway interstitium and has favorable effects on MCT(28–31). Finally, we tested ARINA-1, a nebulized formulation comprised of glutathione (488 mM) and ascorbate (400 mM), as well as sodium bicarbonate (1000 mM). ARINA-1 is currently in development for patients with non-CF bronchiectasis and related chronic inflammatory lung diseases and has also been shown to improve MCT in HBEC models of CF (16, 17). Our rationale for testing Camostat and ivacaftor basolaterally was that this approximates their use as systemic agents. The other compounds tested (PAAG, HA and ARINA 1) are hydrophilic, were used in very low volumes (1 ul) and have been used previously directly on the apical side of airway epithelia. This also mimics their use as inhaled agents.

As an initial screen, we tested the effects of mucoactive agents in primary terminally differentiated epithelial cells at concentrations they are known to affect MCT(16, 26–28). Each compound tested showed some degree of inhibition of SARS-CoV-2 replication in the HBEC (Fig. 2). Ivacaftor was administered to the basolateral compartment and had a moderate effect on SARS-CoV-2 replication. Although some efficacy was observed at 10 µM, its maximum effect was observed at concentrations close to its maximum solubility (30 µM), which is unlikely to be relevant *in vivo* (Fig. 2A). PAAG also showed a moderate effect, with a peak of antiviral activity at 500 µg/ml (in 1 µl volume, applied apically) that decreased at higher concentrations (Fig. 2B). However, both ivacaftor and PAAG showed an increase in cytotoxicity measured by LDH release at the concentrations required to exhibit an effect (Fig. S1, data supplement), which made it difficult to deconvolute whether the observed reduction in viral load was due to the antiviral effects of augmented MCT or reduced replication through cytotoxicity. Due to an inability to rigorously assess these agents under these conditions, they were deprioritized from further evaluation. HA exhibited higher antiviral activity, reaching more than 90% inhibition at a concentration of 0.7% (1 µl, apical) (Fig. 2C). Notably, ARINA-1 exhibited the most efficacious effects on viral replication in epithelial cells, achieving greater than 99% inhibition at the highest concentrations tested. (Fig. 1D). Neither HA nor ARINA-1 caused cytotoxicity at their tested maximum concentrations (0.7% and 100%, respectively, Fig. S1, data supplement). Particularly, high-molecular-weight HA has been evaluated in clinical trials for its potential protection against progression of COVID-19-induced respiratory failure; however, the outcomes of this study have not yet been published(32). On the other hand, ARINA-1 has not been evaluated for its potential benefits in COVID-19 disease. Therefore, due to its novelty, anti-SARS-CoV-2 effectiveness *in vitro*, and emerging clinical safety profile, we further focused our studies on ARINA-1 as a mucoactive agent with potential anti-SARS-CoV-2 activity in respiratory epithelia and evaluated the mechanistic link to ciliary function.

**Fig. 2.**
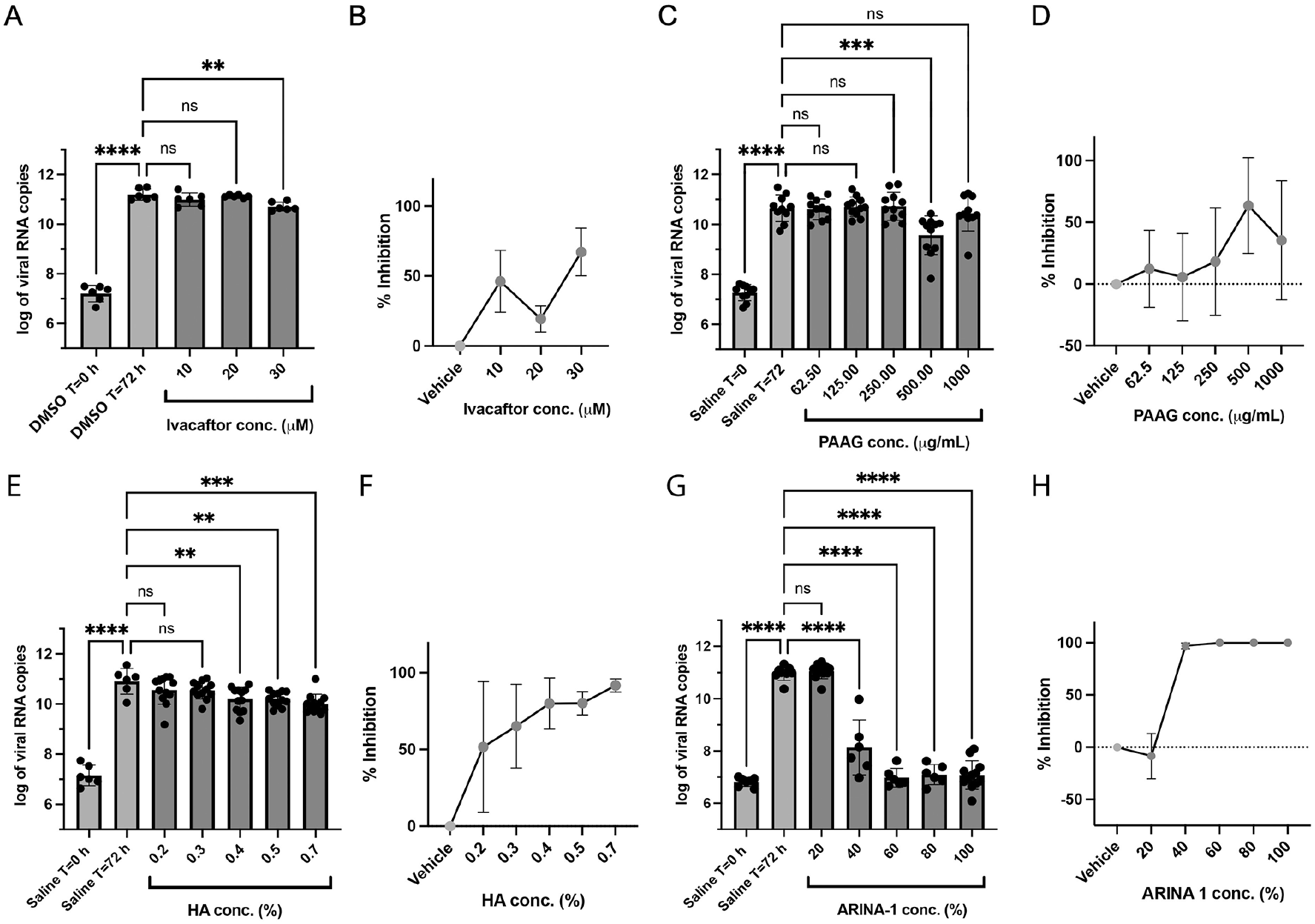
Anti-SARS-CoV-2 activity of the mucoactive agents tested in well-differentiated primary HBE cells. A) Effect on the viral copy number measured by the RT-qPCR caused by ivacaftor at the concentrations shown on the graph. B) Data from graph in A) were converted to percent of viral inhibition. C-D), E-F) and G-H) As in (A-B), but using the compounds PAAG, HA and ARINA-1, respectively. DMSO was the vehicle used for ivacaftor (hydrophobic compound) and saline for PAAG, HA and ARINA-1 (hydrophilic compounds). All experiments were performed at least in duplicate independent assays, each with at least three transwell filter replicates per condition. Treatments were compared using ordinary one-way ANOVA statistical analysis. Hydrophobic compounds (added basolaterally) and hydrophilic (added apically) are shown in green and red, respectively. For each compound, each independent experiment was done with primary HBEC from a different donor.

### ARINA-1 inhibits the shedding of infectious virus by primary HBEC

To ascertain that ARINA-1-caused reduction of SARS-CoV-2 RNA amounts (measured by RT-qPCR) correlated with the blocking of viral replication, we quantified the amount of infectious virus shed by the primary HBEC cells using a median tissue culture infection dose (TCID_50_) determination assay. For this, we washed the apical side of the HBEC to collect virus from ARINA-1-treated or untreated samples and used serial dilutions of the recovered virus to infect Vero E6 cells for its quantification. Thus, we compared the amounts of virus shed by ARINA-1-treated primary HBEC with the mock treated cells (saline) and showed that ARINA-1 blocked the production of infectious virus by two orders of magnitude in HBEC when compared to saline-treated controls (TCID50=1 vs 131-fold dilution, respectively, Fig. 3). This result indicated that the amounts of viral RNA measured by RT-qPCR correlated with the amounts of infectious virus in our HBEC/ALI-based antiviral assay and further validated it as a method for the search of anti-SARS-CoV-2 drugs.

**Fig. 3.**
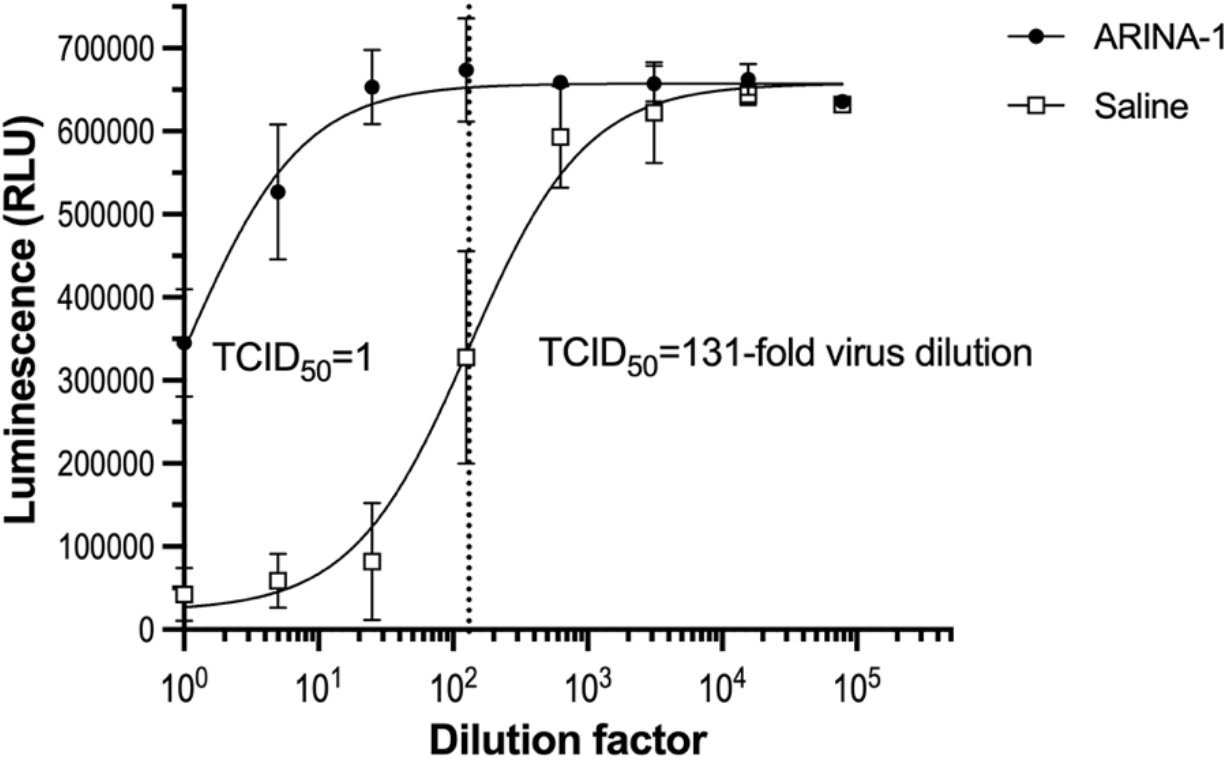
ARINA-1 inhibits the production of infectious virus by primary HBEC. Quantification of SARS-COV-2 virus using a luminescence TCID_50_ assay showed that ARINA-1-treated cells produced two order of magnitude less infectious virus that the saline-treated controls (TCID_50_=1 vs 131-fold dilution, respectively). Values are averages and SD of three independent experiments, each using primary HBEC from a different donor. For each experiment, every dilution of the virus was assessed in sextuplicate using Vero E6 cells as indicators of cytotoxicity using a luminescence assay that measures ATP as a proxy for cell viability.

### ARINA-1 protects HBEC from SARS-CoV-2-driven cytopathogenicity

It has been well documented that SARS-CoV-2 causes extensive plaque-like cytopathic effects in HBECs, including cell fusion, apoptosis, destruction of epithelium integrity, cilium shrinkage, and granular formation on cilia (7, 33, 34). In this study, we observed that well-differentiated primary HBECs from two different donors treated with the vehicle (saline) and infected with SARS-CoV-2 showed the characteristic cytopathic effects caused by this virus (Fig. 4B, F) that were not seen in mock-infected (saline only) controls (Fig. 4A, E). However, the same cells treated with ARINA-1 were protected from the cytopathic effects caused by SARS-CoV-2. Specifically, we observed that ARINA-1 protected the ciliated cells from cilia shrinkage and loss and SARS-CoV-2-induced apoptosis (Fig. 4D, H) Cells treated with ARINA-1 that were infected appeared morphologically similar compared to control cells that were uninfected and treated with ARINA-1 (Fig. 4C, G). In addition, the sections of the mock-treated epithelial layer that showed more cell damage and apoptosis corresponded with higher infection, which was demonstrated by higher immunostaining for a specific viral marker, the S glycoprotein of SARS-CoV-2 (Fig. 4J). However, ARINA-1-treated cells did not show any specific S glycoprotein immunostaining, which strongly indicated that cells were protected from SARS-CoV-2 infection (Fig. 4L) in contrast to the controls (Fig. 4K), suggesting augmented MCT was diminishing cell entry.

**Fig. 4.**
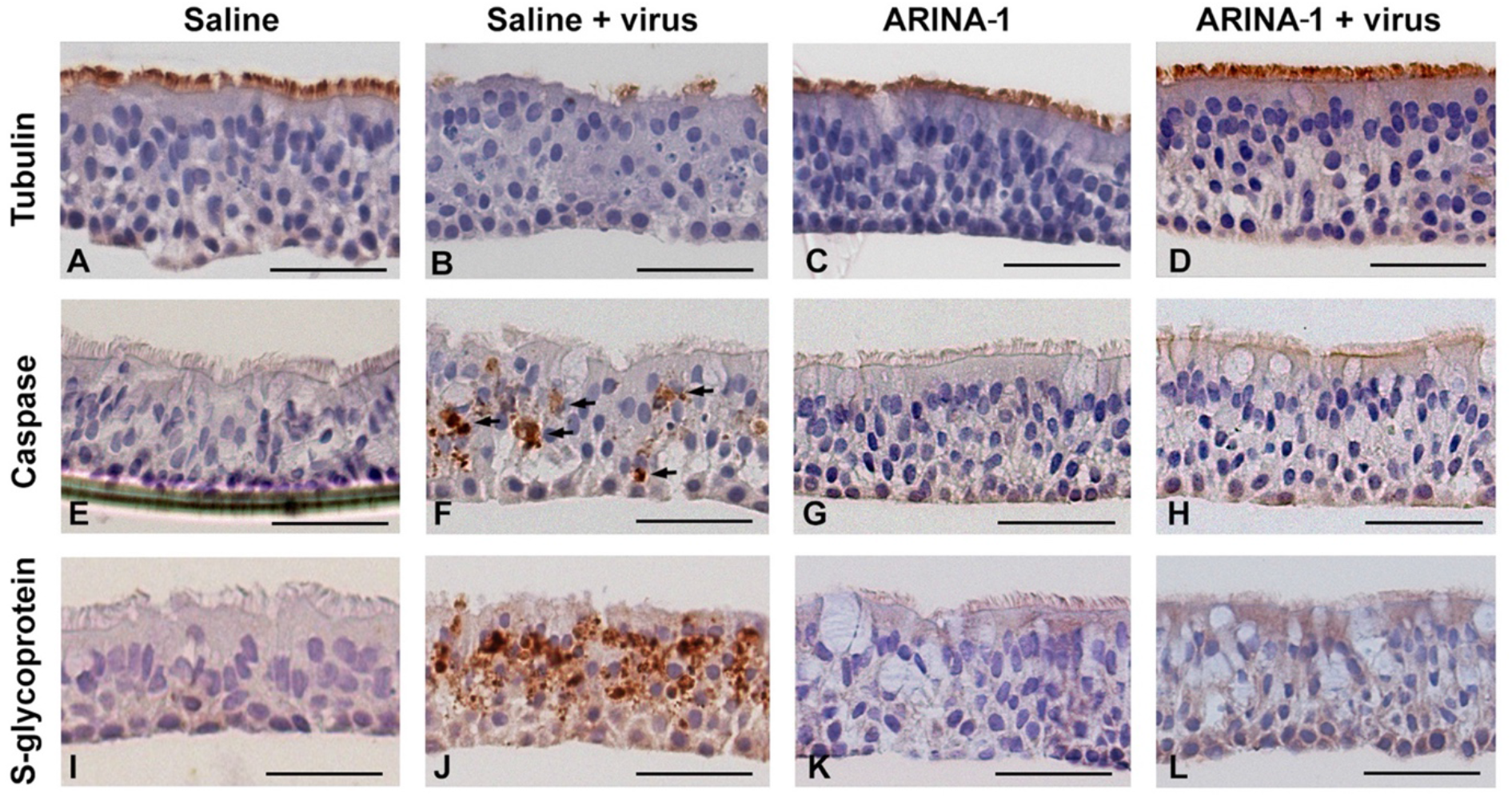
Histopathology studies showed that ARINA-1 protects primary HBE cells from SARS-CoV-2-mediated cytopathology. A-L) Representative photo micrographs of HBEC cross-sections with the immunohistochemistry and treatments. Each row corresponds to the immunohistochemistry using the antibody against the cell marker shown at the left, and each column corresponds to the treatment shown above the upper pictures. SARS-CoV-2 caused cilia shortening and loss in saline-treated cells (panel B). However, ARINA-1 protected cilia from damage (panel D). Similarly, the virus induced significant apoptosis in the mock-treated HBE cells (panel F, apoptotic cells indicated with arrows), which was not observed in those treated with ARINA-1 (panel H). In addition, mock-treated cells showed significant immunostaining using an antibody against the viral S glycoprotein (panel J), which again was not observed in the ARINA-1 treated cells (panel L). Scale bars represent 100 µm. Fully differentiated primary HBEC from two donors were used for the histopathology study. N ≥ 10 pictures per donor and condition were taken and analyzed.

### ARINA-1 induces a supernormal MCT in HBEC and protects from cilia shortening

Previous studies have demonstrated ARINA-1 has beneficial effects on MCT in CF bronchial epithelial cells due to the effects of ascorbic acid, glutathione, and sodium bicarbonate(16). To establish if ARINA-1 beneficially impacts MCT in normal epithelia in the context of SARS-CoV-2 infection, we used micro-optical coherence tomography (µOCT) imaging. µOCT imaging of SARS-CoV-2 infected HBECs with and without treatment of ARINA-1 allowed functional interrogation of the MCT apparatus at the microscopic level to confirm the activation of MCT and the antiviral mechanism of action of ARINA-1. Measurement of cilia beating frequency (CBF) by µOCT showed that in ARINA-1-treated cells, even after infection with SARS-CoV-2, cilia CBF was significantly higher than in the saline-treated controls (Fig. 5A). Correspondingly, MCT was significantly augmented in ARINA-1-treated cells compared to those treated with saline (Fig. 5B), even in SARS-CoV-2 infected cells compared to mock controls (Fig. 5C to F), which indicates that the drug ameliorated the deleterious effect of SARS-CoV-2 on mucociliary function.

**Fig. 5.**
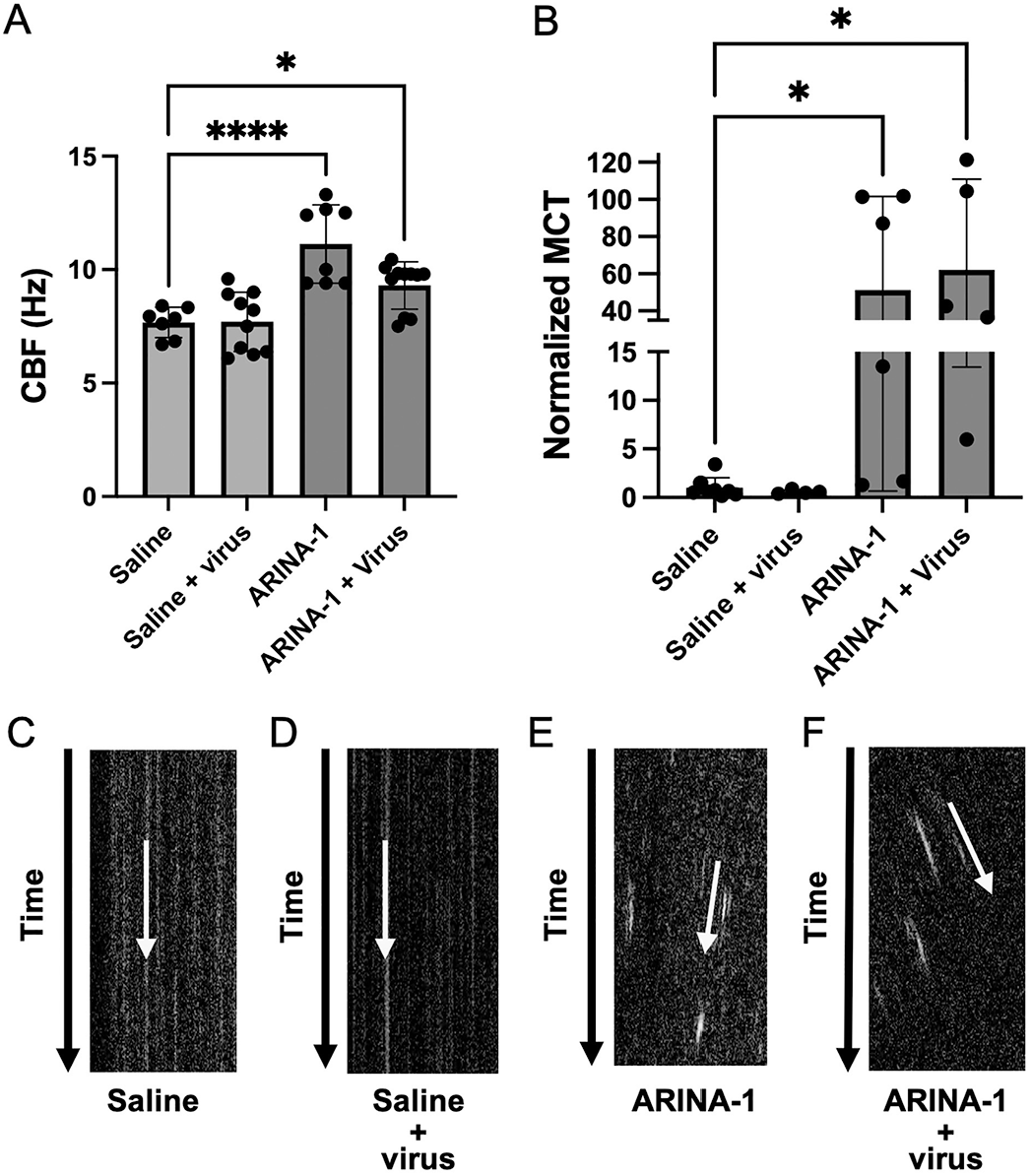
ARINA-1 induces a hypernormal MCT in well-differentiated primary HBEC. A) and B) Cilia beating frequency (CBF) and mucociliary transport (MCT) measurements using µOCT, respectively, in SARS-CoV-2-infected or uninfected, ARINA-1-treated or untreated HBEC Comparisons were performed using an ordinary one-way ANOVA. MCT values were normalized against the MCT average of saline treated baseline controls. C-F) Resliced images of µOCT captured videos (videos S1 to S4 of data supplement) in which the slope of the diagonal streak (yellow arrow) indicates the vectorial transport of mucus particles over time. This allows visualization of MCT rate on still images, in which higher slope angles with respect to time vectors are indicative of faster MCT rates.

The reduction in cilia length, i.e. cilia shrinkage, observed in histology preparations was quantified following the algorithm described in the supplementary *Materials and Methods* section using Image J software. A significant reduction in cilia length was observed in SARS-CoV-2-infected compared to uninfected HBECs (Fig. S2). In contrast, when ARINA-1 was administered 1 hour prior to infecting the cells, cilia shrinkage was not observed (Fig. S2). Together, these results indicated beneficial effects on ciliary structure and function that imparted improved MCT.

### ARINA-1 blocks SARS-CoV-2 replication even when administered after viral infection

Given that ARINA-1 significantly reduced SARS-CoV-2 infection of HBECs when administered one hour prior to introduction of virus, we next determined whether ARINA-1 treatment of cells after initiation of infection would also confer benefit by limiting the spread of infection between cells and ultimately ameliorate total viral load. To test this, we infected HBEC for 1 hour and then washed off excess viral particles, limiting infection of the epithelial cells to endogenous virus. We then treated the cells with ARINA-1 at 3 and 24 hours after infection onset and assessed viral load at 72 hours after exposure to the virus. We observed that ARINA-1 added 3-and 24-hours post-infection both significantly reduced the viral load, albeit to a lower extent than when the cells were pre-treated with the drug immediately before infection (Fig. S3A, data supplement), suggesting its beneficial effect on viral replication could be conferred through diminishing cell-to-cell spread through apical transmission.

### The combination of buffered antioxidants provides antiviral effect of ARINA-1

Given ARINA-1 is a fixed dose combination of ascorbic acid, glutathione, and bicarbonate, we were interested in the contribution of each of these components to the observed antiviral activity, as we have examined previously in its effects on CF mucus viscoelasticity(16).To evaluate this, we tested the antiviral activity of ascorbic acid plus bicarbonate, glutathione plus bicarbonate, and bicarbonate alone, each at the same concentrations in ARINA-1. Bicarbonate was required to buffer glutathione and ascorbic acid because glutathione and ascorbic acid alone have been shown to be cytotoxic due to the low pH of each antioxidant when used alone (16). We found that both ascorbic acid plus bicarbonate and glutathione plus bicarbonate each have a significant antiviral effect in respiratory epithelial cells, and the three-drug combination was most effective. However, bicarbonate alone did not show significant antiviral activity (Fig. S3B, data supplement). These results suggest that the antioxidant components in ARINA-1, ascorbic acid and glutathione, are each important to conferring protection to respiratory epithelial cells and have previously been shown to confer improved MCT in epithelial cells when used in combination with bicarbonate (16). However, a contribution by bicarbonate, beyond its acid buffering capacity such as its indirect influence on MCT (16, 35, 36), cannot be fully discarded because glutathione and ascorbic acid cannot be tested without bicarbonate.

### ARINA-1 is not directly deleterious to SARS-CoV-2 virions

To assess if ARINA-1 affected SARS-CoV-2 virion directly versus augmenting epithelial function, we tested the direct antiviral activity on the SARS-CoV-2 viral particles. We exposed a suspension of the SARS-CoV-2 virus to ARINA-1 and then removed ARINA-1 by ultrafiltration using a pore size that allows only ARINA-1 components to be removed but not the virus particles (Fig. 6A). The virus particles exposed to ARINA-1 were as infective as those exposed to saline solution in Vero E6 cells (Fig. 6B) and in HBEC grown in an ALI (Fig. 6C). These data demonstrate that ARINA-1 is not deleterious to SARS-CoV-2 virion. Instead, these data support that improved epithelial function is the principal mechanism of its antiviral effects.

**Fig. 6.**
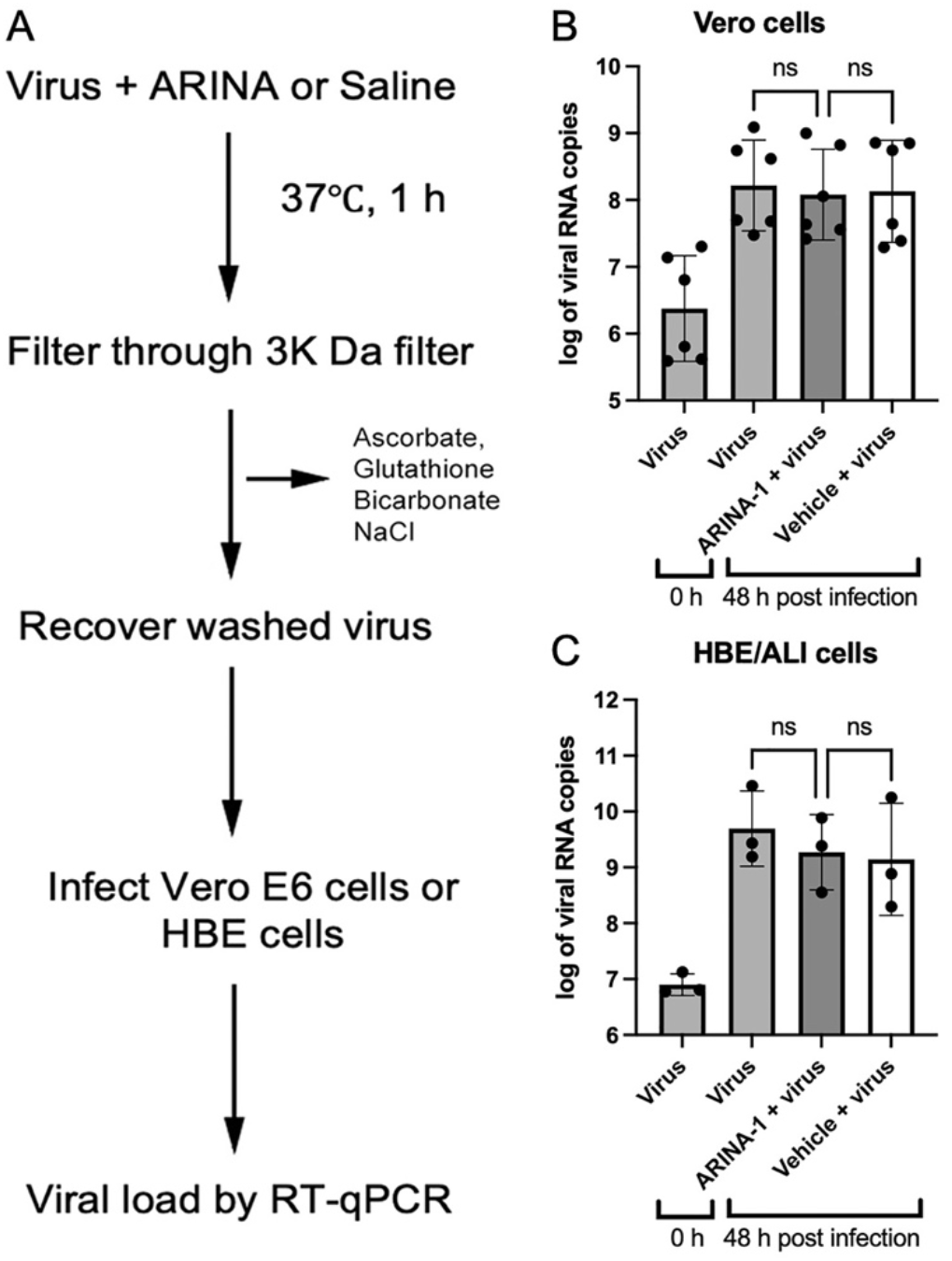
ARINA-1 has no direct antiviral effect on SARS-CoV-2 virus. A) Flow diagram showing the procedure followed to test the direct antiviral activity of ARINA-1 on the virus. Briefly, a suspension of SARS-CoV-2 virus was exposed to ARINA-1 and then filtered through a 3,000 Da pore size membrane to remove the ARINA-1 components. The ARINA-1-free virus suspension was used to infect Vero E6 or HBE/ALI cells, and the viral load determined after 48 h post infection. Virus particles were treated in parallel with saline as control. B) Viral load of the ARINA-1-treated virus compared to the saline treated virus measured in Vero E6 cells or in C) the HBEC/ALI assay. RNA copy numbers were logarithmically transformed and compared using ordinary one-way ANOVA statistical analysis. ns: non-significant.

### MCT accounts for the anti-SARS-CoV-2 protection conferred by ARINA-1

Given the efficacy of ARINA-1 in blocking SARS-CoV-2 infection, we further investigated if its antiviral activity can be explained by MCT improvement alone or if there may be another ARINA-1-triggered cellular mechanism of protection involved or interference with the viral entry and post-entry pathaways. To answer this question, we devised a viral replication assay in which MCT was blocked in the primary HBECs using an inhibitor of cilia beating or by using undifferentiated HBEC where cilia have not developed yet, and then evaluated if ARINA-1 still conferred protection from infection. For the first approach, MCT was blocked using BAPTA/AM, a membrane-permeable selective calcium chelator that inhibits cilia motility(37). Since calcium is required for cilia motility, sequestering it by BAPTA/AM immobilizes the cilium beating. In the presence of BAPTA/AM (20 µM), ARINA-1 did not exhibit antiviral activity (Fig. 7A), and BAPTA/AM was not cytotoxic at the concentration used (Fig. S1, data supplement). For the second approach and to discard possible off-target effects of BAPTA/AM, we used immortalized 16HBE14o-cells and primary HBECs grown at ALI for only one week, before mucociliary differentiation is achieved, so cilia were not present in either of these conditions and MCT was therefore absent (38, 39). Each cell line was treated with ARINA-1 or saline and then infected with SARS-CoV-2 (Fig. 7B and C). Under these conditions, ARINA-1 did not inhibit SARS-CoV-2 infection, indicating MCT must be present to exhibit its inhibitory properties on SARS-CoV-2 replication.

**Fig. 7.**
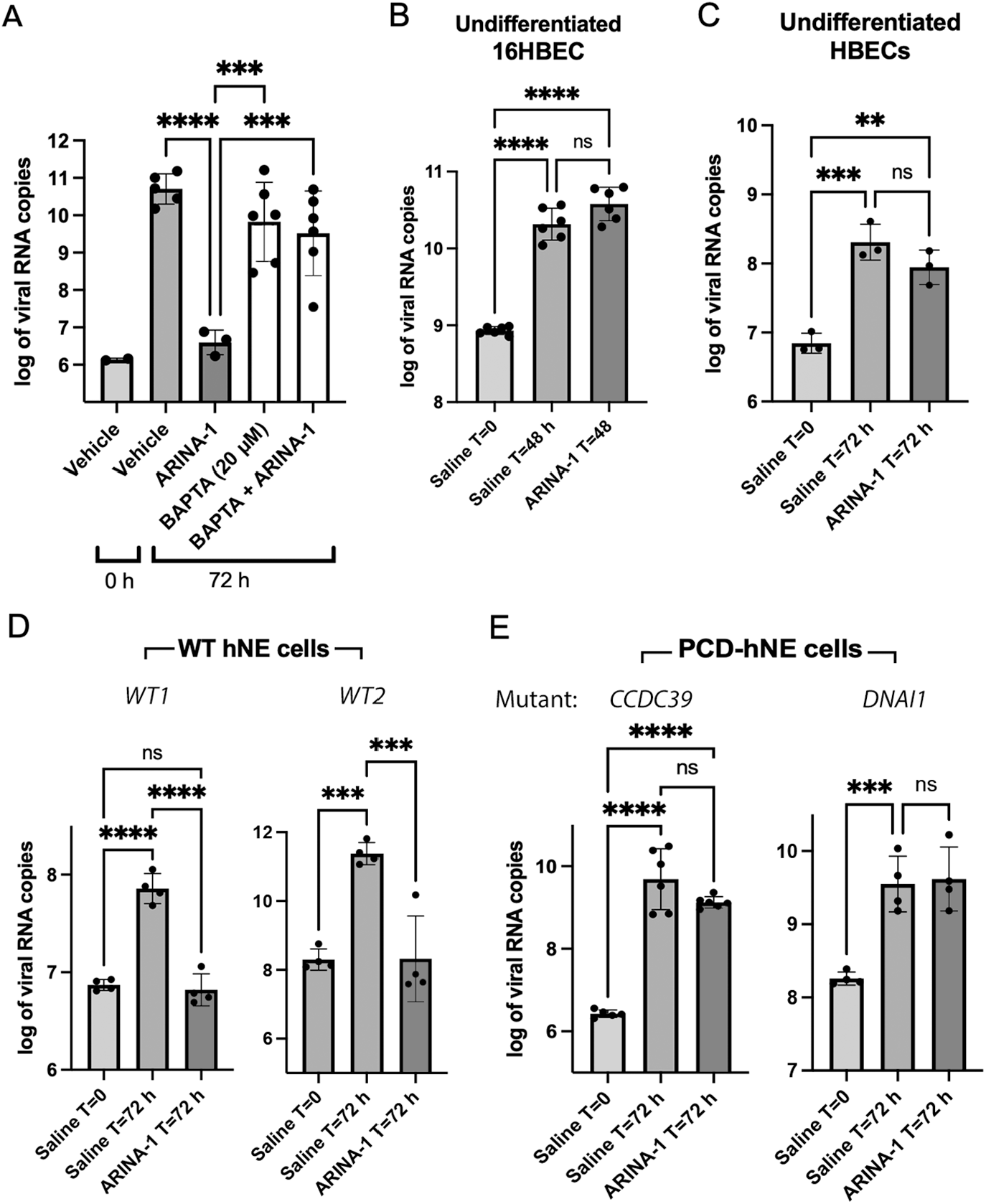
ARINA-1 is not protective when cilia beating is inhibited with BAPTA/AM or cilia are not active or present in cells. A) Viral loads in HBEC treated with ARINA-1, BAPTA/AM or ARINA-1 plus BAPTA/AM. Two independent experiments with 3 technical replicates for BAPTA/AM and BAPTA/AM plus ARINA-1 were performed. B) Viral loads of undifferentiated 16HBE and C) primary HBEC, which lack cilia, were treated with ARINA-1 or saline as control. Two independent experiments with 3 technical replicates for 16HBE and one experiment with 3 replicates for HBEC were performed. D) Viral loads of ALI differentiated hNE cells from two WT donors and E) from two PCD suffering human donors with mutations in genes *CCDC39* and *DNAI1*, which encode proteins essential for the assembly of dynein arm complexes and for dynein protein itself, respectively, treated with ARINA-1 or saline as control. Two experiments with at least two technical replicates for PCD-hNE cells and one experiment with four replicates for each WT-hNE cell donor were performed. Data were logarithmically transformed and compared using ordinary one-way ANOVA.

As a final test, we evaluated ARINA-1 in human nasal epithelial cells (hNEs) from two patients affected by primary ciliary dyskinesia (PCD) syndrome, a genetic condition characterized by defective cilia expression or function (40). Similar to the HBEC, hNE cells from healthy donors (WT-hNE cells) and from the PCD donors (PCD-hNE cells) were terminally differentiated at ALI, and then the effect on SARS-CoV-2 replication assessed. ARINA-1 did not protect against SARS-CoV-2 infection in the PCD-hNE cells; however, it remained inhibitory in the WT-hNE cells (Fig. 7D and E, respectively). Taken together, these results indicate that ciliary motility is required for ARINA-1-dependent antiviral protection and that MCT is the primary mechanism of action of the antiviral effect of ARINA-1 seen in the ALI-primary HBEC infection model.

### The oxidative state of the cells influences antiviral protection

Proper cilia functioning requires a balanced redox environment, which is maintained by redox regulatory proteins and thioredoxin domain-containing proteins (41). Dynein ATPases, which are the ciliary protein motors, as well as several cilia-localized proteins that modulate dynein activity (e.g. Protein Kinase A, Protein Kinase C, and Protein Phosphatase 1), are sensitive to the local redox environment within each cilium (41). As noted, the active constituents of ARINA-1 are ascorbic acid and glutathione, with bicarbonate acting as a buffering agent. Since both ascorbic acid and glutathione are known for their antioxidant capacity, we next investigated whether the antioxidant aspect of the molecules is responsible for the improvement of the MCT and consequent antiviral activity. We hypothesized that decreasing the endogenous production of reactive oxygen species (ROS) by HBECs can simulate anti-SARS-CoV-2 protection conferred by ARINA-1. To reduce endogenous ROS production, we inhibited xanthine oxidase. Xanthine oxidase catalyzes the oxidation of hypoxanthine to xanthine and can further catalyze the oxidation of xanthine to uric acid, generating ROS in the process. Hypoxanthine is derived from the catabolism of ATP via AMP. Thus, energy-demanding activities that consume ATP, such as cilia beating, generate ROS through the xanthine oxidase pathway (42–44). Inhibition of xanthine oxidase with allopurinol significantly protected the HBEC from SARS-CoV-2 infection (Fig. 8A).

**Fig. 8.**
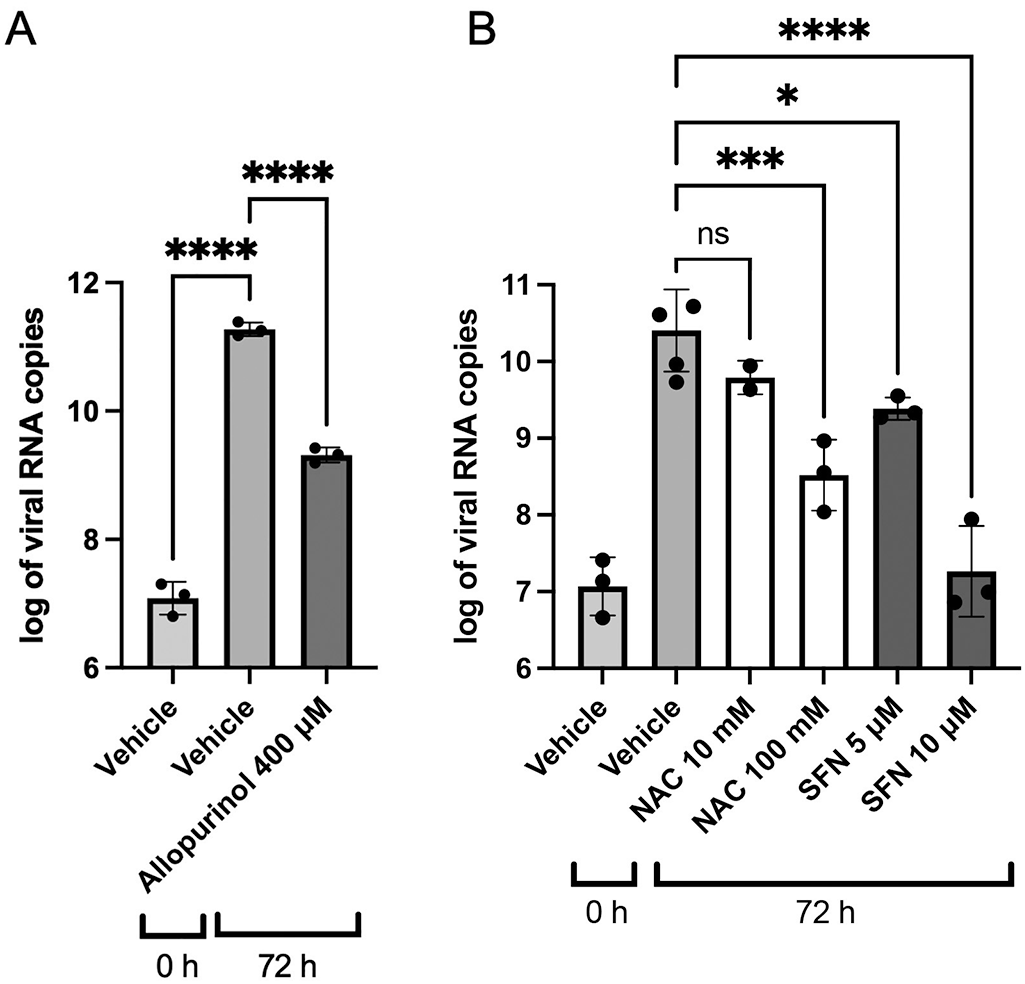
A proper redox state of the cell is essential for the antiviral protection conferred by mucociliary transport. A) Viral load in the presence of allopurinol (400 µM) compared to vehicle (DMSO) in the HBEC/ALI assay. B) Viral load in the presence of the antioxidant agents N-acetylcysteine (NAC) and sulforaphane (SFN) at the specified concentrations.

We also tested if antioxidants other than ascorbic acid and glutathione could also confer antiviral protection. We used N-Acetyl-L-cysteine (NAC) and sulforaphane (SFN) to test this hypothesis (45). As expected, both compounds caused a reduction in the viral load when tested in the HBEC/ALI antiviral assay (Fig. 8B), and neither compound showed cytotoxicity at the higher concentrations tested (400 µM, 100 mM, and 10 µM, respectively (Fig. S1B, data supplement). These results suggest that antioxidants may benefit epithelial cell resistance to SARS-CoV-2 by imposing a redox state favorable to MCT and may explain, at least in part, the beneficial effects of ARINA-1.

## Discussion/Conclusion

Using a model of terminally differentiated HBEC grown in an ALI, we demonstrated that mucoactive agents have the capacity to block SARS-CoV-2 infection in vitro. Of the five agents tested, each selected based on their capacity to augment MCT in differentiated respiratory epithelia, HA, Camostat mesylate and ARINA-1 demonstrated the most significant SARS-CoV-2 inhibition (Fig. 1 and 2). We further studied ARINA-1 based on its safety profile and novelty as a potential anti-COVID drug and showed that its inhibitory effect on SARS-CoV-2 replications was dependent on ciliary function. These results provide an important advance: targeting MCT may block SARS-CoV-2 replication, in addition to its potential effects protecting against SARS-CoV-2 induced mucus stasis and secondary infections (46–48).

As an initial control, we evaluated camostat mesylate, which provided almost complete inhibition of viral replication at concentrations higher than 16.6 uM (Fig. 1). This drug has been previously shown to have partial anti-SARS activity in Vero E6 cells by inhibiting TMPRSS2 (22). However, we observed complete inhibition of SARS-CoV-2 replication in our HBEC/ALI model, which may be explained by its additional mucoactive properties via ENaC inhibition. The mucoactive action of camostat could not be evaluated in Vero E6 cells (monkey kidney epithelial cells), which are not specialized airway epithelial cells.

These results suggested evaluation of other mucoactive agents with mechanisms of action that do not alter viral replication per se was warranted. Among these, ARINA-1, a formulation comprised of a combination of glutathione, bicarbonate and ascorbate, that is in development for chronic inflammatory airways diseases such as non-CF bronchiectasis, exhibited a complete inhibition of SARS-CoV-2 replication (Fig. 2). ARINA-1 is known to augment MCT through its properties on mucus viscoelasticity and ciliary function (16). Likewise, HA showed dose-dependent antiviral activity with a half-maximal inhibitory concentration of around 0.2% and achieved up to 92% inhibition at 0.7%, which is within the range of HA concentrations used in clinical studies for respiratory diseases (31). In contrast to the other agents tested, ivacaftor and PAAG showed some SARS-CoV-2 inhibition in respiratory epithelia but were associated with cytotoxicity at the conditions required to impart benefit (Fig. 2E), which made discerning what portion of the reduced viral load is due to the direct antiviral activity of the drugs or to cytotoxicity challenging.

Given the potential for use of ARINA-1 in SARS-CoV-2 infection, we further characterized this drug. We showed that ARINA-1-mediated inhibition of SARS-CoV-2 replication correlated with full protection of the HBECs from the characteristic SARS-CoV-2 cytopathic effect that was observed in the saline-treated cells (Fig. 4). Using µOCT, we observed that ARINA-1 increased the MCT to a level significantly higher than the normal rate in HBEC (Fig. 5B). Although this experiment did not show an effect of the virus on cilia beating or mucociliary clearance, this does not mean there is not a deleterious effect of the virus on these parameters, but that the technique did not have the power to resolve differences in cilia beating or MCT values between infected and uninfected cells of the mock controls (baseline values using saline instead of ARINA-1, Fig. 5). Unlike our work using similar techniques in hamsters (49), the reduction of MCT by virus in primary HBE cells was not evident. We think this is because the MCT values of HBEC mock controls (i.e. those treated with saline and not infected) measured by µOCT were very low at baseline, and thus, to measure an additional reduction in MCT by the virus is difficult to detect. In our experience, HBEC from some donors show low MCT values by micro-OCT at baseline, perhaps due to dehydrating conditions of incubator growth, despite the attempt to humidify them in that scenario; this was intrinsically more difficuilt to control in the BSL3 environment.

We propose a model for the ARINA-1 protection mechanism in which the hyper-normal MCT phenotype caused by the drug hinders virus attachment to the cell receptor, preventing the cell from viral infection. In our *in vitro* assay conditions, the viral particles are constantly moving in the airway surface liquid at a rate that does not allow their access to the cell surface and engagement with the host cell receptor. Even when ARINA-1 was administered 24 h after viral infection, the drug showed significant antiviral protection (Fig. S3A, data supplement). SARS-CoV-2 infection of HBECs has been shown to occur in a plaque-like manner (7, 33, 34). The virus undergoes cycles of replication in which the virus particles produced in one plaque are released and disperse to infect distant cells, forming new plaques or infection foci. It is possible that ARINA-1 administered post-infection can prevent the spread of infection from initial foci of infection to new cells in the vicinity by improving MCT in the uninfected cells surrounding the plaques.

As none of the components of ARINA-1 are known to have a direct antiviral effect, our initial results suggested that the inhibition of SARS-CoV-2 infection was likely due to the mucoactive properties of ARINA-1^27^. To confirm, we studied if factors other than MCT could account for the antiviral effect observed in the HBEC model grown at ALI. First, we tested if ARINA-1 had a direct antiviral activity against SARS-CoV-2 virion. After exposing SARS-CoV-2 viral particles to ARINA-1 for an hour, the virions remained fully infectious to Vero or HBE cells (Fig. 6B and C), indicating that ARiNA-1 has no deleterious effect on the virus itself. Second, we investigated if ARINA-1 interferes with viral entry or post-entry mechanisms or activates another cellular mechanism that accounts for some of its antiviral activity. After removing the contribution of MCT from the antiviral assays by blocking the cilia movement with BAPTA/AM (Fig. 7A), using undifferentiated HBEC that are devoid of cilia (Fig. 7B and C), and by evaluating its effects in hNE cells from PCD-suffering donors, where cilia are present but dysfunctional (Fig. 7E), the cells were not protected from SARS-CoV-2 infection in the presence of ARINA-1 as occurred in cells from WT hNE donors (Fig. 7D). Furthermore, ARINA-1 did not alter the cellular entry or post-entry pathways of SARS-CoV-2 infection since the virus was able to strongly infect HBE and PCD-hNE cells that did not exhibit ciliary expression or function in the presence of ARINA-1 (Fig. 7). Taken together, these results indicate that ARINA-1-mediated improvement of MCT is the primary mechanism of ARINA-1 antiviral activity in the airway epithelial cells tested.

It is known that SARS-CoV-2 and other viral and bacterial infections cause oxidative stress in infected cells (50–52). In addition, there is a redox regulation of cilia motility in the airway’s epithelium through various mechanisms (41). Therefore, we hypothesize that SARS-CoV-2 increases ROS to a level that is detrimental to cilia movement, which prevents viral clearance and favors the attachment of newly formed virus. We also propose that the antioxidant capacity of ARINA-1 reduces the excess ROS induced by SARS-CoV-2 to a level that not only neutralizes the deleterious effect of the virus on cilia movement, but also improves the antioxidant capacity to supranormal levels with consequential improvements in cilia motility and MCT. Our studies on allopurinol inhibition support this hypothesis. We demonstrated that allopurinol inhibition of endogenous ROS generation by xanthine oxidase blocked SARS-CoV-2 infection (Fig. 8), Additionally, the use of the other antioxidants NAC and SFN also blocked SARS-CoV-2 infection (Fig. 8). It is plausible that antioxidant agents act on unknown components, in addition to the known ones, of the cilium machinery and its regulatory network, each one providing an additive effect on cilia movement that is beneficial for MCC.

In summary, we demonstrate that augmenting MCT can have beneficial effects on SARS-CoV-2 replication in respiratory epithelia, and that several agents that are clinically available or are in late-stage development can confer these beneficial properties. These studies emphasize the importance of mucociliary clearance as a barrier against SARS-CoV-2 replication and severity and demonstrates the significant potential impact that improved ciliary function in the airway epithelia may impart, deserving further exploration in COVID-19 and potentially other respiratory infectious diseases.

## Acknowledgement (optional)

The authors thank Carolyn Durham, PhD and Dan Copeland from Renovion, Inc. for their critical review of our manuscript and the patients who donated cells for research. The authors also thank Dezhi Wang and John Ness from the UAB Pathology Core Research Laboratory/Department of Pathology for tissue processing, slicing and immunostaining.

## Statement of Ethics

The UAB Institutional Review Board approved the use of human cells and tissues (IRB-080625002), which were obtained after written informed consent from donor subjects.

## Author Contributions

J.C-G. and S.M.R. conception and design of the research. J.C-G., C.F.P., M.M., L.T., R.J., Q.L., J.E.P.L., S.H., performed experiments. J.C-G, G.M.S., E.O., H.K., S.M.R. and K.H. analyzed data. J.C-G and S.M.R. interpreted the results of experiments. J.C-G., C.F.P., Q.L., J.E.P.L., S.H. and S.M.R. prepared figures. J.C-G. and S.M.R. drafted the manuscript. J. C-G., C.F.P., J.E.P.L. and S.M.R. edited the manuscript. All authors approved the final version of the manuscript.

## Data Availability Statement

The data that support the findings of this study will be made available upon reasonable request from the corresponding author.

## SUPPLEMENTAL DATA

Supplemental Figs. S1–S3 and Supplemental videos S1-S4:

https://figshare.com/s/a81607ab32aab4045561

