## Supplementary data for "Mucociliary Clearance Augmenting Drugs Block SARS-Cov-2 Replication in Human Airway Epithelial Cells"

ONLINE DATA SUPPLEMENT

### SUPPLEMENTARY MATERIALS AND METHODS

#### Procurement and growth of human primary airway epithelial cells.

The UAB Institutional Review Board approved the use of human cells and tissues. Primary HBEC were derived from lung explants after written informed consent was obtained from donor subjects using methods described previously (1, 2) [see details in data supplement]. Briefly, tissues were debrided immediately after surgical resection, washed twice in Minimum Essential Media with 0.5 mg/ml DTT (Sigma-Aldrich, St. Louis, MO) and 25 U/ml DNAse I (Roche, Basel, Switzerland), and then placed in dissociation media containing MEM, 2.5 U/ml DNAse I, 100 mg/ml ceftazidime, 80 mg/ml tobramycin, 1.25 mg/ml amphotericin B, and 4.4 U/ml pronase (Sigma-Aldrich) for 24–36 h at 4°C. Loosened airway epithelial cells were then expanded using BEGM medium (LONZA, Basel, Switzerland) supplemented with an additional 10 nM all-trans-retinoic acid (Sigma-Aldrich) that was exchanged every 24 h. Following expansion, first or second passage cells at 80–90% confluency were dissociated and seeded at a density of $0.5 \times 10^5$ on Costar® Transwell 24-well filter inserts (cat. # 3470, Corning Inc.) after coating with NIH 3T3 fibroblast conditioned media. Cells were differentiated at ALI for at least five weeks, at least otherwise is specified, using medium PneumaCult™/heparin/hydrocortisone (Stemcell™ Technologies) plus pen/strep antibiotics (exchanged three times a week) before further use. Cells from five different donors were used in this study: WT128, WT148, WT152, WT158 and WT210 cell lines.

Nasal cells from a healthy and a PCD patient [genotype: heterozygous pathogenic CCDC39 c.2586+1G>A (Splice donor) and CCDC39 c.830_831del (p.Thr277Argfs*3)] were obtained from nasal brushes after written informed consent was obtained from donors and grown as co-culture with 3T3 J2 cells. When the cells become 80-90% confluent the cells were detached using 0.05% trypsin. A total of $1.25 \times 10^5$ cells were plated in FNC coated Costar® Transwell 24-well filter inserts (cat. # 3470, Corning Inc.). After three days medium from the apical compartments was removed and differentiation medium (Pneumacult ALI maintenance medium) maintained at the basolateral side only. Medium was changed every other day until terminal differentiation (two and a half to three weeks) and then used for the experiments.

**Culture of 16HBE at the ALI.**

16HBEC14o- were grown using EMEM medium (ATCC, cat. # 30-2003™) supplemented with 10% FBS and were dissociated at 80–90% confluence and seeded at a density of $0.5 \times 10^5$ on Costar® Transwell 24-well filter inserts (cat. # 3470, Corning Inc.) after coating with FNC (Athena ES 0407H) and using EMEM medium. Upon reaching confluence, the medium was removed from the apical side, and the cells were grown at the ALI for no longer than two weeks to avoid differentiation. They were then used in the antiviral assay described below.

for quantitative study of airway functional microanatomy using micro-optical coherence tomography. *PLoS One* 2013;8:.

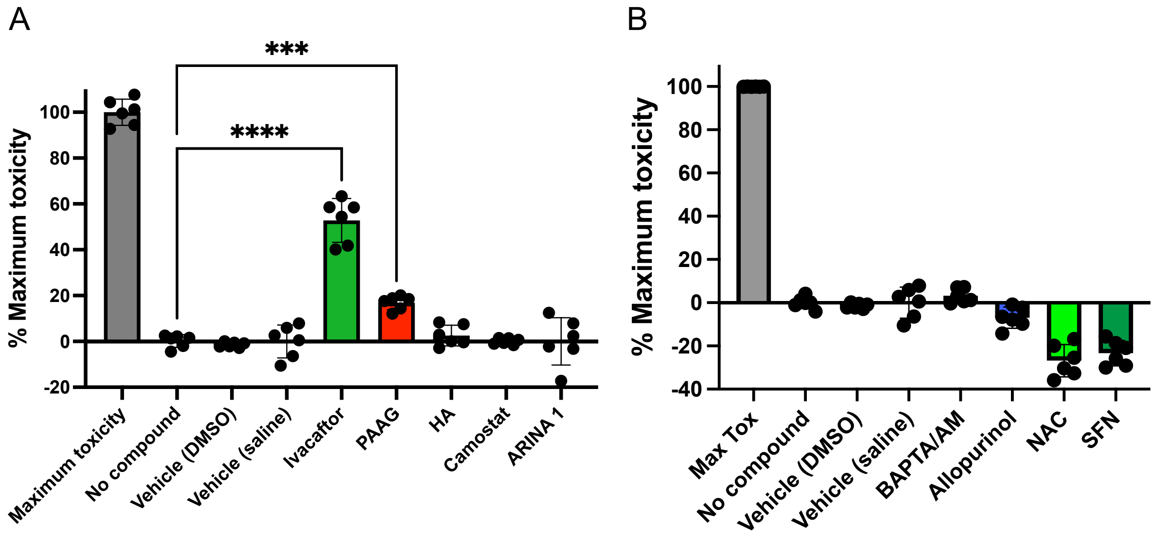


**Figure S1.** Cytotoxicity of the compounds tested in this study A) Cytotoxicity of the mucoactive agents tested B) Cytotoxicity of additional compounds used in the study. The maximum concentration used in the studies was used to evaluate cytotoxicity of each compound (Ivacaftor, 30 μM; PAAG, 1 mg/mL; HA, 0.7%; Camostat, 125 μM; Allopurinol, 400 μM; NAC, 100 mM and SFN, 10 μM). DMSO was the vehicle used hydrophobic compounds (Ivacaftor, Camostat mesylate, BAPTA/AM, Allopurinol and SFN) and saline for hydrophilic compounds (PAAG, HA, ARINA-1 and NAC). All experiments were performed at least in duplicate independent assays, each with three transwell filter replicates per condition. Treatments were compared using ordinary one-way ANOVA statistical analysis. Allopurinol, NAC and SFN showed negative values of toxicity indicating they preserved cell viability for the duration of the assay (72 hours).

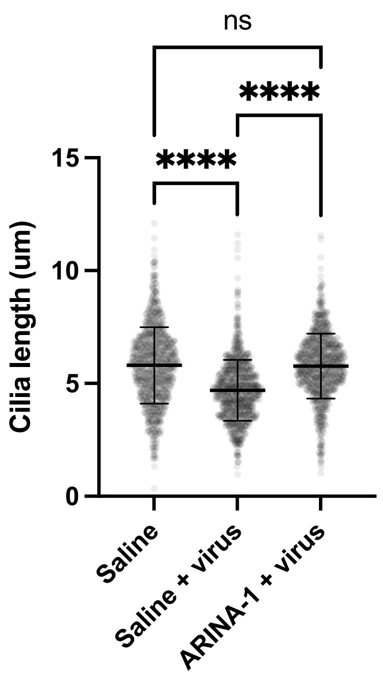


**Figure S2.** ARINA-1 protects from cilia shrinkage. Cilia length comparison of uninfected untreated HBEC with SARS-CoV-2-infected mock-treated or ARINA-1-treated HBEC. Comparisons were done using ordinary one-way ANOVA (N=969 cilia were measured for each condition).

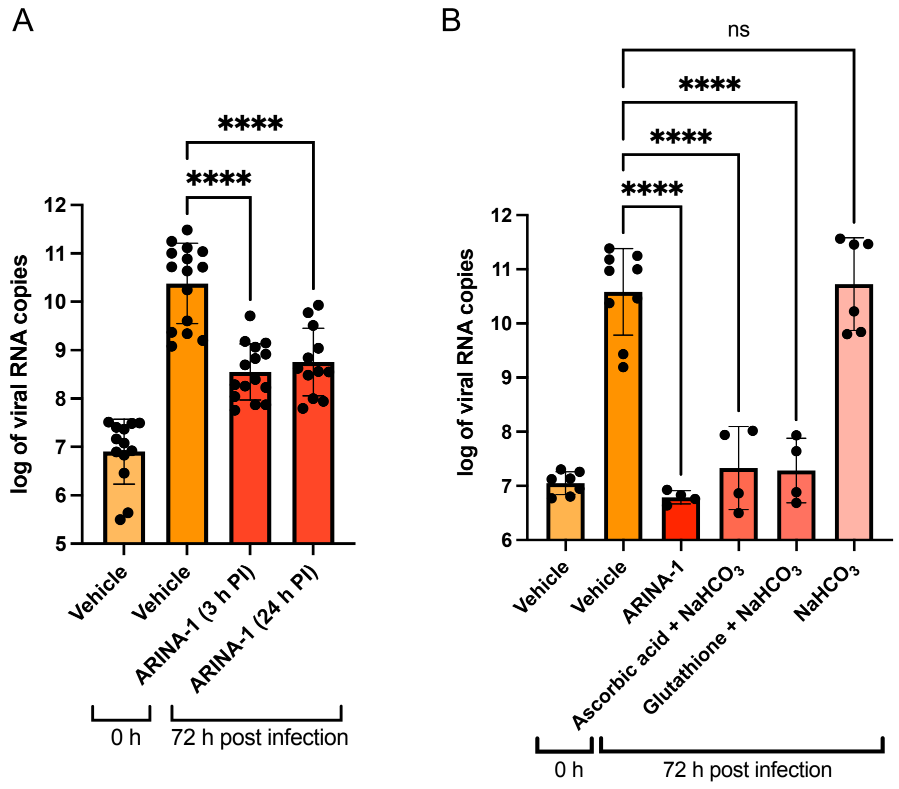


**Figure S3.** ARINA-1 blocks SARS-CoV-2 replication when administered after viral infection and its antiviral components are ascorbic acid and glutathione. A) Viral load (measured 72 hr after virus exposure) of HBEC treated with ARINA-1 (1 µl apical) 3 and 24 hours after exposing cells to virus, compared to vehicle (saline)-treated cells. Three independent experiments were performed with at least 3 technical replicates per condition. B) Antiviral activity of the ARINA-1 components. Because ascorbic acid and glutathione alone are toxic to cells due to their acidities, ascorbic acid plus sodium bicarbonate, glutathione plus sodium bicarbonate, and sodium bicarbonate alone at the same concentrations in ARINA-1 were assessed in the antiviral assay. Two independent experiments were done with at least two technical replicates per condition. For both data sets (A and B) RNA copy numbers were logarithmically transformed and compared using an ordinary one-way ANOVA statistical analysis.
